# Behavioural rhythms of two gammarid species *Echinogammarus marinus* and *Gammarus pulex* under increasing levels of light at night

**DOI:** 10.1101/2024.09.18.613239

**Authors:** Charlotte N. Underwood, Alex T. Ford, Samuel C. Robson, Herman Wijnen

## Abstract

Artificial light at night (ALAN) is proliferating at an alarming rate across the globe, particularly around aquatic habitats. Natural and predictable light cycles dictate much of an individual organism’s life by acting as a major signal for their circadian clock, driving rhythmic behaviours and physiological changes throughout the body. Light cycles also help populations coordinate group behaviour and greatly impact the interspecies dynamics of a community. Research into the ecological impacts of ALAN has highlighted numerous effects on these biological processes, including higher predation rates, impaired growth and development, and diminished reproductive success. Invertebrates play an undeniable role in ecosystem functioning and show robust daily rhythms. As such, it is vital to understand how ALAN may disrupt their behavioural patterns. The aim of this study was to monitor the impacts of increasing levels of light at night (0 lux – 80 lux), as well as constant light and constant darkness, on the behavioural rhythms of the intertidal amphipod, *Echinogammarus marinus*, and the freshwater species, *Gammarus pulex. G. pulex* activity was not strongly synchronised to any of the light at night treatments. *E. marinus*, however, exhibited strong behavioural rhythmicity in diurnal cycles with dark night periods. All the ALAN treatments resulted in a significant decrease in *E. marinus* rhythmicity and overall activity. Moreover, ALAN between 1-50 lux disrupted nocturnality in this species. These results indicate that while some gammarids show some adaptive plasticity when it comes to light pollution, others may experience strong direct impacts on their activity. This may be relevant to individual and population level fitness of vulnerable species in more heavily urbanised areas.

## Introduction

The proliferation of the human population has led to a dramatic increase in artificial light at night (ALAN) around the world. More than 80% of people live under conditions that exceed natural night sky brightness (Falchi et al., 2016; Garstang, 1989; Kyba et al., 2015) and the amount of artificially lit outdoor area is increasing by 2.2% each year (Kyba et al., 2017). The influence of artificial light extends beyond urbanised areas due to skyglow, the phenomena in which artificial light is scattered and reflected back to earth by the atmosphere. Skyglow is capable of masking lunar brightness (Davies et al., 2013b) and has resulted in significant increases in natural light levels over 30 km from major cities (Kyba et al., 2011). As a result, more than 22% of coastlines are impacted by ALAN globally (Davies et al., 2014) and an estimated 35% of marine protected areas around the world experience ALAN (Davies et al., 2016). Although there is currently no large-scale analysis of ALAN in freshwater environments (Jechow and Hölker, 2019), 90% of people live within 10 km of at least one substantial freshwater body such as a lake or river (Kummu et al., 2011), meaning that ALAN has the potential to encroach on aquatic ecosystems both inland and towards the sea.

The widespread transition to broad-spectrum lighting like white LEDs will introduce ecosystems to night-time lighting at shorter wavelengths compared to older technologies (Gaston et al., 2013). This blue light is visually detectable by a wider range of taxa (Davies et al., 2013a; Gaston et al., 2013) and propagates more easily through the atmosphere and through water (Markager and Vincent, 2000; Warrant and Locket, 2004), increasing the scope of light pollution in both terrestrial and aquatic habitats, an effect that carries many ecological implications.

The most thoroughly studied area of chronobiology has undoubtedly been the circadian clock, the biochemical oscillator set by the solar day. Consistent light cycles play a significant role in life’s evolutionary history, shaping the daily rhythms found in nearly all organisms (Bhadra et al., 2017). Light helps establish ecological niches, mediates physiological processes, and influences population dynamics and interspecies relationships (Longcore and Rich, 2004). Numerous studies have highlighted the ecological impacts of ALAN on processes known to be controlled by the circadian clock across a wide range of species, both terrestrial and aquatic (Davies and Smyth, 2018). Many aspects of animal behaviour and physiology have been affected to some degree, with little understanding of the potential cascading effects and long-term consequences for ecosystems. Regardless of whether an animal is positively or negatively phototactic (attracted or repelled by light, respectively), the presence of light at abnormal times can alter their natural behavioural patterns, affecting their ability to forage, find mates, and avoid predation (Bird and Parker, 2014; Luarte et al., 2016; Russo et al., 2019). By providing more favourable conditions for some taxa over others, light pollution can upset the community structures and dynamics that have evolved over millennia (Barker and Cowan, 2018; Davies et al., 2015), meaning that ALAN has consequences for individual fitness, as well as the intraspecies relationships, interspecies competition, and trophic interactions that make up an ecosystem.

Invertebrates are vulnerable to changes in light intensity and spectrum (Davies et al., 2012; Garratt et al., 2019; Holzhauer et al., 2015; Stewart, 2021), and it is crucial to assess how their behaviour and physiology are impacted by ALAN and determine any long-term risks associated with chronic exposure. Amphipods are an order of crustaceans that have colonized virtually all aquatic ecosystems. The intertidal species *Echinogammarus marinus* (Leach, 1815) and the freshwater species *Gammarus pulex* (Linnaeus, 1758), are commonly found throughout Europe (Dick et al., 2005; Karaman and Pinkster, 1977). Despite their significant ecological roles as nutrient recyclers and as a food source for higher trophic levels, there has been limited research into the impacts of light pollution on their behavioural and physiology (Perkin et al., 2014). Their typically negative relationships with light are known to be reversed by parasitism and exposure to certain pharmaceuticals (Bakker et al., 1997; Guler et al., 2015; Guler and Ford, 2010; Perrot-Minnot et al., 2013), which leaves them at a potentially higher risk of predation in polluted waterways that are exposed to light at night. There is also evidence that sex determination is linked to changes in photoperiod in *E. marinus* (Guler et al., 2012), meaning the proliferation of artificial light could have significant consequences for individual health, population dynamics, and distribution. The aims of this research are to examine how increasing levels of ALAN affect the behavioural rhythms, activity levels and nocturnality in these two species of amphipod.

## Methods

### Animal collection and husbandry

Adult *E. marinus*, along with one of their main foods, the macroalgae *Fucus vesiculosus*, were collected by hand during low tide from Lock Lake, Portsmouth, United Kingdom (N 50°47’22.7”, W 1°02’30.6”) between July 2020 and February 2022. *G. pulex* were collected from River Ems, Westbourne, United Kingdom (N 50°51′34.8”, W 0°55′45.8”) using the kick sampling method (Mackey et al., 1984) and 1 mm mesh nets between November 2020 and July 2021. On one occasion, *G. pulex* was collected from a culture at the University of Portsmouth, Institute of Marine Sciences, where the animals were contained in a 18L aquarium with water and organic debris collected from River Ems and then aerated with an aquarium pump and airstone. Half of the water in the aquarium was changed every three days and replaced with fresh river water. The aquarium was kept at 18°C under a 12h:12h light/dark cycle. Individuals from this aquarium were randomly chosen using an aquarium net. After collection, both *E. marinus* and *G. pulex* were transported to the University of Southampton, Life Sciences Building.

At the University of Southampton, each species was maintained in its own 5L capacity tank in an incubator set to 10°C and 70% humidity. For *E. marinus*, artificial seawater was prepared using Instant Ocean (Aquarium Systems, Sarrebourg, FR) and distilled water to attain a salinity of 33 ppt. *F. vesiculosus* was added to their tank to provide food and shelter. Water from River Ems was primarily used in the *G. pulex* tank, with Evian mineral water substituted in when river water was unavailable. Organic debris and sediment from River Ems were included in the tank to provide food and shelter. In both cases, the animals were allowed to feed ad libitum. The tanks were aerated using an aquarium pump and airstone, and then covered with a lid to limit evaporation. Half of the water in the tanks was changed twice per week and replaced with new water. Wastewater was filtered through a 70 µm mesh to prevent animal loss. The incubator was equipped with a programmable light (Fluval Plant Spectrum LED) that allowed for gradual transitions from light to dark and vice versa over three hours. Amphipods were left in laboratory conditions for two weeks under a 12h:12h light/dark (LD) cycle prior to beginning any behavioural assays to allow for acclimation and for any tidal rhythms to dissipate in *E. marinus*. Daylight consisted of all LEDs on the Fluval light turned up to the maximum setting (2750 lux; 6500K).

### Behavioural assays

To assess vertical movement in the water column, behavioural assays were conducted using Locomotor Activity Monitors (LAM25, TriKinetics Inc., Waltham, MA, USA), which count activity via the breaking of infrared beams (1 beam break = 1 count). An approximately equal number of sexually mature males and non-gravid females (n_total_= 32) were assigned to cylindrical vials (dim. 25 mm x 95 mm) filled with ∼40 ml of either artificial seawater or river water, depending on the species of amphipod. The artificial seawater was first mixed with *E. marinus* tank water (2:1 ratio) to reduce its purity and minimise shock to the animals upon transfer to the vials. Prior to this, both seawater and river water were aerated with an airstone for 24 hours to ensure adequate oxygenation. During the assays, they were given a piece of food, *F. vesiculosus* for *E. marinus*, or organic debris for *G. pulex*. The vials were plugged with a cotton bung to minimise water loss from evaporation and were placed vertically in the LAM25. The activity monitors, along with a *Drosophila* Environment Monitor (DEnM, TriKinetics Inc., Waltham, MA, USA), were placed in an incubator set to 10°C and 70% humidity. All monitors were connected to a Dell Latitude D620 laptop located outside the incubator running DAMSystem3 *Data Collection Software* (TriKinetics Inc., Waltham, MA, USA) on Windows XP. The monitors were left for eight days to ensure a full seven days of uninterrupted data collection.

Both species were exposed to eight light regimes (Table 1). Natural night sky brightness ranges from 0.0001 lux under a cloudy night to 0.3 lux under a full moon (Gaston et al., 2013; Kyba et al., 2017). The 12h:12h light/dark treatment (LD) was thus used as a control to simulate natural conditions. The first ALAN treatment (LA01) was selected to determine the lowest light level necessary to mask lunar light and elicit behavioural changes. 1 lux was the lowest light level the environmental monitor could detect. The remaining four ALAN treatments (LA05, LA30, LA50, LA80) were selected based on the range of light measurements taken from nearshore marine environments exposed to artificial light pollution (Davies et al., 2015; Jelassi et al., 2014; Lynn et al., 2021). Daylight was the same as the acclimation period (2750 lux; 6500K). A constant light (LL) and a constant dark (DD) treatment were included to monitor free-running circadian behaviour (Table 1).

**Table 1.** Light conditions for behavioural assays. Day phase length included the three-hour transitions at lights on/off, except under constant conditions (LL and DD).

| Regime | Phase length (hours) |  | Max. light intensity (lux) |  |
| --- | --- | --- | --- | --- |
|  | Day | Night | Day | Night |
| LD | 12 | 12 | 2750 | 0 |
| LA01 | 12 | 12 | 2750 | 1 |
| LA05 | 12 | 12 | 2750 | 5 |
| LA30 | 12 | 12 | 2750 | 30 |
| LA50 | 12 | 12 | 2750 | 50 |
| LA80 | 12 | 12 | 2750 | 80 |
| LL | 24 | 0 | 2750 | 2750 |
| DD | 0 | 24 | 0 | 0 |

Cold white light (15000K) was used for the light at night treatments. We chose a white light treatment as one of the main concerns regarding the frequent use of white LEDs for street lighting around the world is that they will introduce more blue light into nightscapes compared to older technologies (Falchi et al., 2016; Gaston et al., 2013; GVR, 2021). These short wavelengths have a comparatively stronger impact on behaviour and physiology (Geoffroy et al., 2021; Guo et al., 2011; Jackson and Moore, 2019; Li et al., 2022). Czarnecka et al. (2022) found that freshwater amphipods from both light-naïve and urban habitats were both repelled by blue light.

### Activity analysis

After the eight-day period, the data was collected via DAMFileScan software (TriKinetics Inc., Waltham, MA, USA). The activity data was grouped into five-minute bins and then analysed and plotted using ClockLab Analysis 6 (Actimetrics, Wilmette, IL, USA). The data was normalised by dividing the individual results by the group mean and then averaging. This data was used to measure the rhythmicity, period length, total activity, and nocturnality of the gammarids. Rhythmicity was analysed using *X*^*2*^-periodograms from the circadian range (15 – 35 h) with 0.01 significance levels. To determine if the individuals showed rhythmic behaviour, the relative rhythmic power (RRP) was calculated by taking the ratio of the peak amplitude divided by the significance threshold (Goda et al., 2011).

Behaviour was considered arrhythmic if the RRP was less than 1 (<1); weakly rhythmic if it was greater than or equal to 1 but less than 1.5 (≥1–1.49); and strongly rhythmic if it was greater than or equal to 1.5 (≥1.5). Only the period length of rhythmic animals was considered. Nocturnality was calculated as the proportion of activity counts during the designated night phase (ZT12:00 – ZT23:59), or the subjective night phase if conditions were constant (i.e. LL, DD; Table 1), divided by the total activity counts.

### Statistical analysis

Only data from individuals that survived the full eight days and were sufficiently active (total counts ≥ 200) was used in the statistical analyses. The data from one channel in the constant light treatment was also discarded as it was discovered at the end of the assay that the vial contained two *G. pulex* individuals. Due to high mortality of *E. marinus* under constant darkness, an additional assay was conducted to verify the results and enhance the statistical power of the analysis. *E. marinus* and *G. pulex* data was analysed separately.

The percentage of animals that survived the assays, the percentage of surviving individuals that remained active, and the ratios of strongly rhythmic, weakly rhythmic, and arrhythmic individuals were each compared using Fisher’s exact test. Normality of the behavioural parameters (total activity, RRP, period length of rhythmic individuals, and nocturnality) were tested using the Shapiro-Wilk test. The impacts of the night-time light treatment on those parameters were then analysed using non-parametric Kruskal-Wallis Rank tests followed by pairwise Mann-Whitney U-tests with Bonferroni corrections to adjust for multiple comparisons. For each light regime, the survival and proportion of active males and females were compared using Fisher’s exact test. Mann-Whitney U-tests were also used to assess differences in total activity, RRP, period length, and nocturnality between the sexes. All statistical analyses were conducted using the software R (V4.0.2, R Core Team, 2020).

## Results

### Differences in female and male responses to light at night

Initial separate analyses for male and female individuals (Table S1) uncovered little behavioural difference between male and female animals. This was the basis for pooling male and female data in the following analyses to obtain increased statistical power. The few significant sex-specific differences detected are outlined here for transparency.

*E. marinus* males had lower survival and activity rates under LA80. Alternatively, while females had 100% survival under constant light (LL), the proportion of active females was only 50%. Their total counts were also significantly lower (Figure S1C), and they became arrhythmic in contrast to the males, which were strongly rhythmic (Table S1; Figure S1A; Figure S2). A Spearman’s Rank Correlation test revealed a strong positive relationship between total counts and RRP under LL (rho = 0.676; p < 0.001), meaning the primary sex difference for *E. marinus* in LL might be that of reduced activity in females.

The proportion of active female *G. pulex* was less than the males under LD and LA30. Females were significantly less active under LA01 and constant darkness (DD; Figure S1D & E). They were also arrhythmic under LA30, whereas the males remained weakly rhythmic (Table S1; Figure S1B; Figure S2).

### Percentage of living and active individuals

100% of *E. marinus* remained alive and active under the LD treatment. Comparatively, survival was significantly lower under four of the treatments (LA05, LA50, LA80, DD), with constant darkness yielding the highest mortality (Table 2). The proportion of surviving individuals that remained active was only significantly reduced in constant light (LL).

**Table 2.**
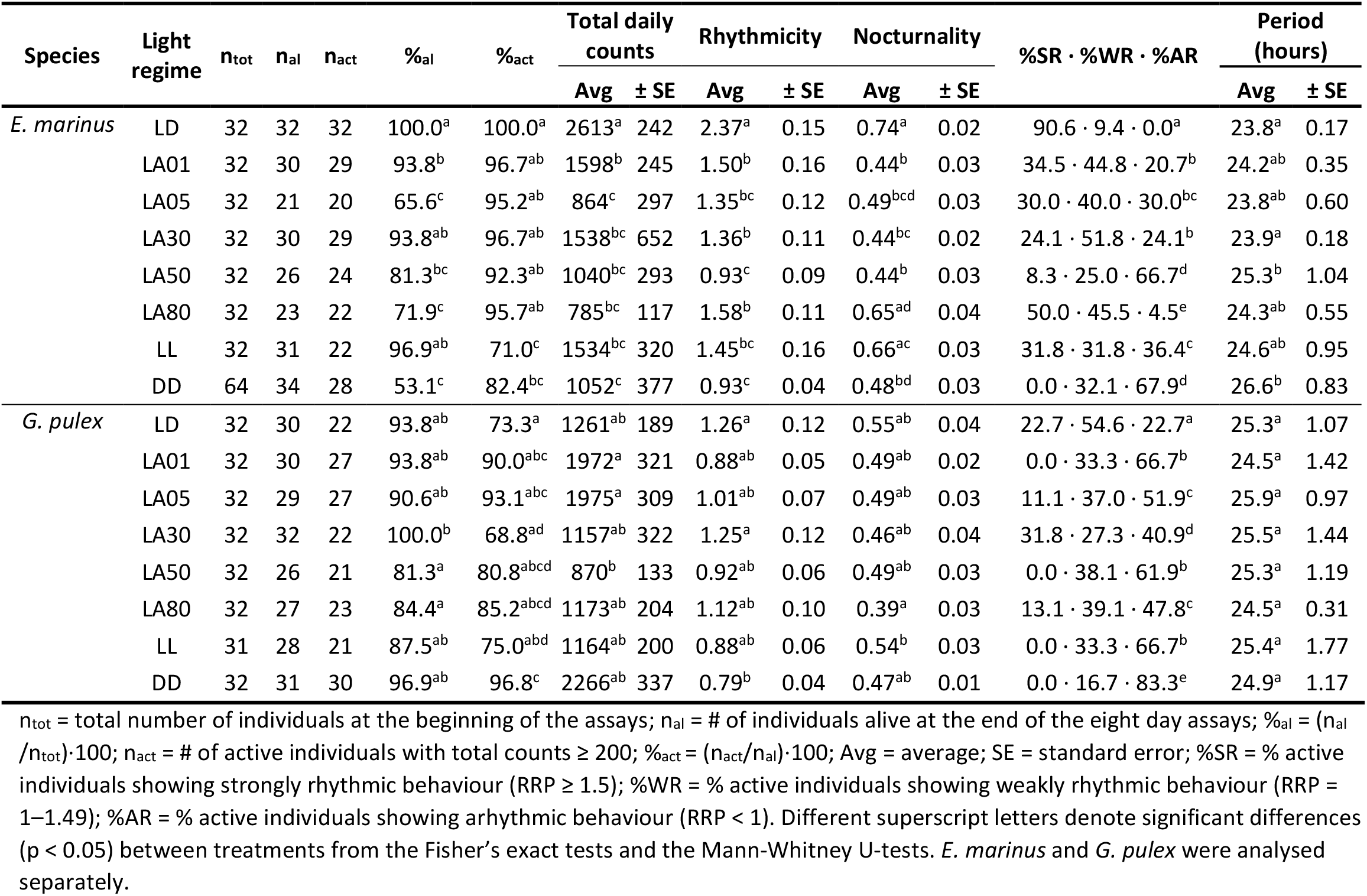
Behavioural parameters of *E. marinus* and *G. pulex* activity under varying light/dark cycles.

Overall, the average survival was higher for *G. pulex* (91.4%) compared to *E. marinus* (81.7%), but *E. marinus* had a higher percentage of active individuals (Table 2). The percentages of surviving *G. pulex* was not significantly affected by the light treatments, but the proportion of active animals was significantly higher under DD.

### Activity rhythms

Under a 12h:12h LD regime, *E. marinus* displayed activity that was strongly rhythmic (RRP = 2.37 ± 0.15; Table 2), with an approximate 24-hour cycle (period = 23.8 ± 0.17 hours; Table 2) and two activity peaks, one at ZT0 and a larger one right after lights off at ZT12. These peaks were either lost or dampened under the other light treatments, although they generally still occurred around the on/off transitions (Figure 1). Total activity significantly decreased (p <0.05; Figure 2A) and rhythms were significantly weakened (p <0.05; Figure 2C) under all other conditions relative to LD. The mean power of behavioural rhythmicity remained strongly rhythmic (SR) under LA80 but was weakly rhythmic (WR) under most of the other light treatments, only becoming completely arrhythmic (AR) under LA50. When rhythmicity was maintained, however, none of the ALAN treatments significantly changed the period length of 24 hours; only constant darkness (DD) saw a significant lengthening of the circadian period (Figure 2E). The ratio between SR:WR:AR individuals significantly shifted under all light conditions compared to LD, moving towards more WR and AR individuals overall (Table 2). Under the LD regime *E. marinus* behaviour was predominantly nocturnal (0.74 ± 0.02; Table 2), with the proportion of night phase activity decreasing in the other light treatments, becoming significantly lower under all but LA80 and LL (Figure 2G).

**Figure 1.**
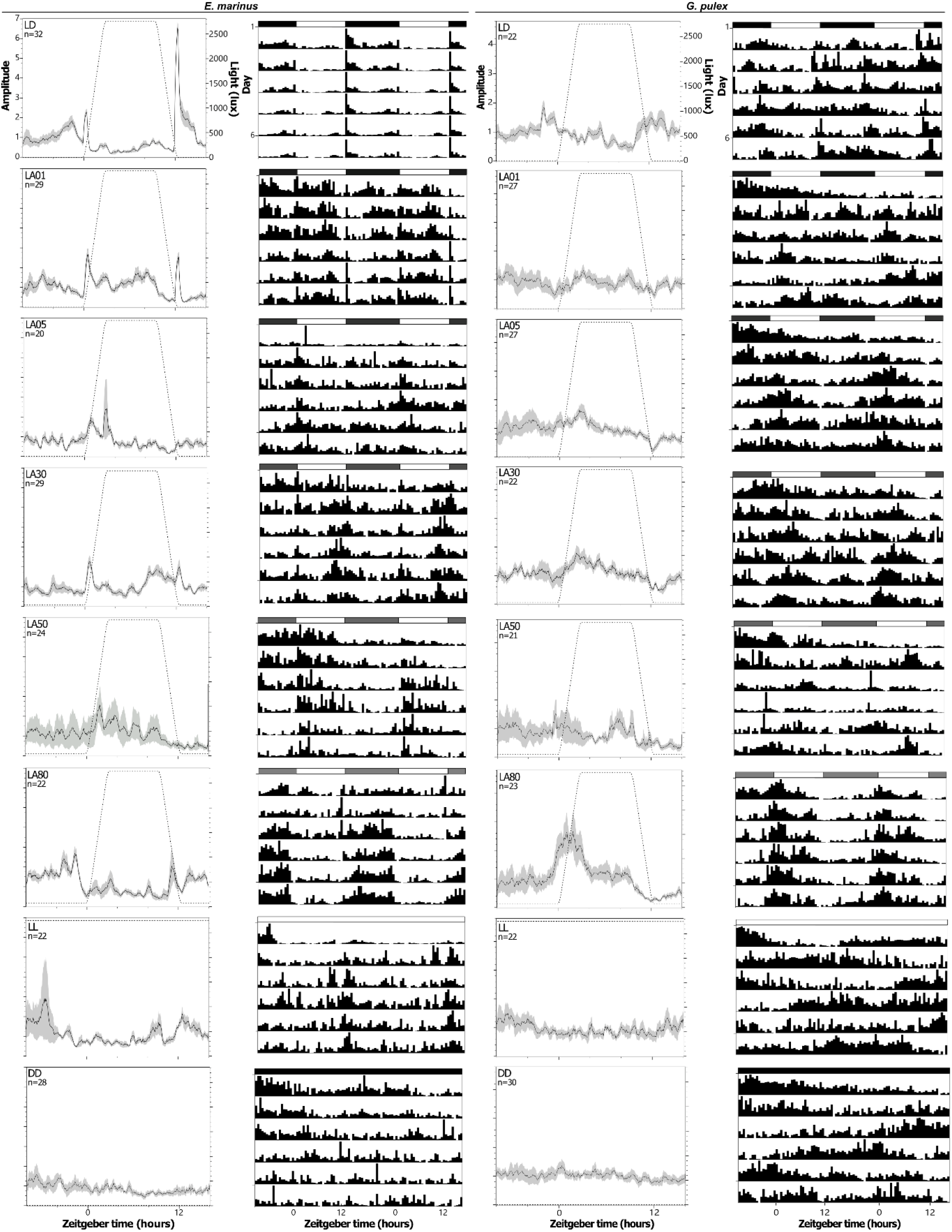
Average activity profiles (columns 1 & 3) and double-plotted actograms (col. 2 & 4) of *E. marinus* (col. 1 & 2) and *G. pulex* (col. 3 & 4). Activity profiles display average activity levels (black line) with standard deviation (grey areas), along with the light levels (dotted lines) over 24 hours. Actograms display the average, normalised behaviour across the 7-day assays, with each row showing two 24-hour cycles. Bars above actograms denote light levels; on/off transitions are not shown. Y axes are the same within columns. Light conditions outlined in Table 1.

**Figure 2.**
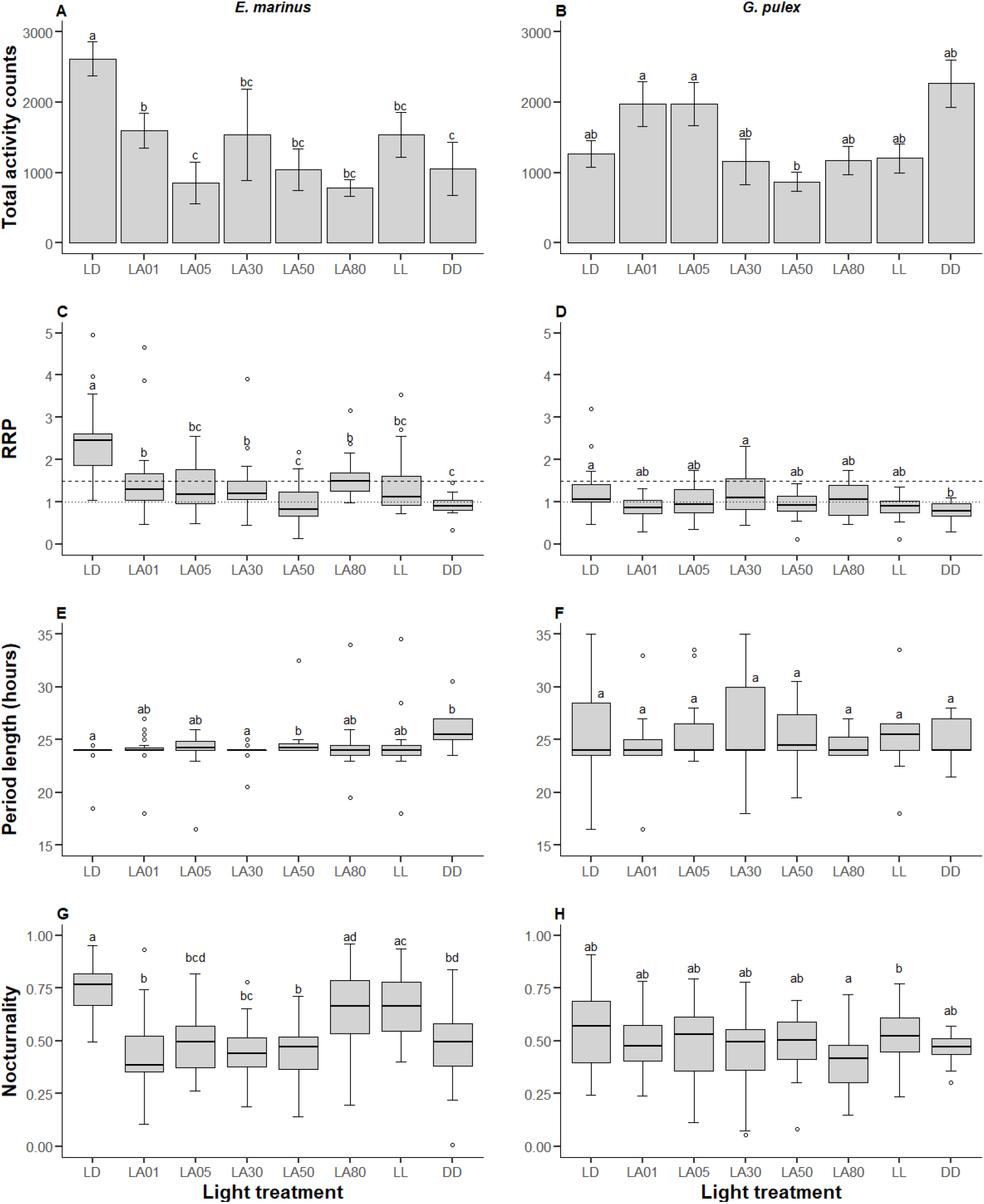
Locomotor activity of *E. marinus* (left column) and *G. pulex* (right column). (A-B) Barplots with standard error bars of total amount of activity counts logged during the seven-day assay period. (C-D) Boxplots of relative rhythmic power (RRP), which shows how strongly the activity repeated over a set period (e.g., 24 hours). Dotted line = 1 (rhythmic); Dashed line = 1.5 (strongly rhythmic). (E-F) Boxplots of periodicity, which shows the length of the phase of repeated behaviour in individuals with RRP ≥ 1. (G-H) Boxplots of nocturnality, which shows the proportion of activity counts during the subjective night phase. Different lowercase letters indicate significant differences (p < 0.05) between treatments. Light conditions outlined in Table 1.

Under a 12h:12h LD regime, *G. pulex* displayed activity that was weakly rhythmic (RRP = 1.26 ± 0.12; Table 2) over an approximate 24-hour cycle (period = 25.3 ± 1.07, Table 2), with two activity peaks, the highest around ZT22 and another at ZT12 (Figure 1). These peaks were lost or dampened under all other light treatments, except LA80, although activity was generally higher around lights on. Total activity increased under the low ALAN treatments (LA01 and LA05) and constant darkness (DD) but decreased at higher levels of light at night (LA30, LA50, LA80, and LL; Figure 2B). Activity became arrhythmic (RRP <1) under all but three light conditions (LA05, LA30, and LA80) but rhythmicity was only significantly lower under constant darkness (Figure 2D). Although the period lengths showed high variance, it was not affected by the light treatments (Figure 2F). The ratio between SR:WR:AR individuals was significantly altered under all the light treatments (Table 2). *G. pulex* did not show a strong preference for either the day or the subjective night phase under any of the light conditions, although nocturnality was lowest under LA80 (Figure 2H).

## Discussion

The two amphipod species from this study demonstrated notably different patterns of activity under the LD regime, which was meant to simulate a natural 24-hour light/dark cycle. While both species were quite sensitive to even low light intensities, as illustrated by the peaks in activity around the light-dark transitions, *E. marinus’* behaviour was strongly rhythmic over a 24-hour period compared to the weak rhythmicity of *G. pulex*. Furthermore, *E. marinus* activity was clearly upregulated in dark phase, whereas *G. pulex* activity was more evenly distributed throughout the day. *E. marinus* had a strong reaction to the different light treatments, lowering their activity, becoming less rhythmic, and in most cases, shifting away from strict nocturnality. Alternatively, *G. pulex* behaviour was not significantly impacted by any of the light treatments, except constant darkness which significantly reduced rhythmicity.

Under a few light conditions, both species showed reduced activity or rhythmicity in females compared to males. Notably, there were no significant sex-specific differences in the behaviour of *E. marinus* exposed to LD or ALAN cycles. On average, males are larger than females (Conlan, 1991), and there is some evidence that females have shorter lifespans, likely due to the high cost of reproduction (Marques and Nogueira, 1991). However, there is no conclusive evidence that these differences would elicit the contrasting behaviours between males and females witnessed in this study, particularly as they were not consistent across the light treatments.

The disparities in overall *E. marinus* and *G. pulex* activity and their responses to light at night may be attributed to differing foraging strategies and predator types. *E. marinus* live amongst one of their primary food sources, *F. vesiculosus* (Martins et al., 2014), and are mostly predated upon by birds during low tide (Múrias et al., 1996), and fish during high tide (Alexander et al., 2013; Beermann et al., 2018). Minimising movement when exposed to light might be an adaptive response to avoid detection by predators. Meanwhile, *G. pulex* drift through their habitat in search of leaf litter (Flecker, 1992), avoiding fish predation through continual movement (Bakker et al., 1997). Although presumed to be nocturnal (Flecker, 1992; Müller, 1963), our results suggest that *G. pulex* does not necessarily favour the cover of darkness. Similar differences between the two species have previously been observed in regards to their phototactic behaviour, with *E. marinus* actively avoiding lit areas more than *G. pulex* (Kohler et al., 2018). Given that we found no significant differences between any of the ALAN treatments in terms of total activity counts, rhythmicity, period length, or nocturnality, it is possible *G. pulex* could be more resilient to the impacts of light at night. We cannot exclude that the increased levels of activity under LA01 and LA05 are reflective of increased foraging efficiency. Nevertheless, ALAN conditions might also increase vulnerability to visual predation (Yurk and Trites, 2000). Previous studies have found that light pollution has no prolonged impact on *G. pulex* drift rates (Perkin et al., 2014), while exposure to predator cues does not elicit long-term reductions in activity and feeding rates (Åbjörnsson et al., 2000; Szokoli et al., 2015). Increased feeding observed in other freshwater gammarids exposed to light at night is hypothesised to be due to changes in metabolic rates as a result of light-induced stress (Czarnecka et al., 2021; Manríquez et al., 2019), meaning if *G. pulex* activity levels can recover after prolonged exposure to ALAN, it may actually be driven by a higher energetic demand. The prioritisation of foraging over predator avoidance regardless of environmental conditions has also been observed in dogwhelks, which were more likely to forage under ALAN regardless of the presence of a predator cue (Underwood et al., 2017). Further investigations into *G. pulex* activity and feeding rates under lower light levels are necessary to discern whether ALAN exposure is beneficial or detrimental to their individual fitness.

*E. marinus* experienced significantly lower survivorship under three of the ALAN treatments. While exposure to ALAN can elicit increased rates of oxygen consumption (Manríquez et al., 2019), which would have depleted the limited oxygen in the vials, there was not consistently high mortality across all treatments with night-time light exposure. Under constant darkness, the unusually high mortality could be partially attributed to oxygen levels. Each vial contained a piece of *F. vesiculosus* to act as shelter and food for the animals throughout the assays. If the seaweed continued to photosynthesise under the other light conditions, this would have provided additional oxygen throughout the experiments, but this would not have been the case under constant darkness. Nevertheless, the low survival rates cannot be conclusively connected to the light conditions without further investigations, which should include monitoring oxygen levels throughout the assays.

Any level of light at night significantly weakened *E. marinus* behavioural rhythms. There was also a noticeable decrease in activity across all light at night treatments. These results are consistent with previous work examining *E. marinus* escape responses to light (Kohler et al., 2018), as well as the impacts of ALAN on talitrid amphipods whose activity was either arrhythmic (Lynn et al., 2021) or reduced (Bohli-Abderrazek et al., 2018; Jelassi et al., 2014; Luarte et al., 2016). While ALAN did not lead to any changes in period length, *E. marinus* did appear to transition from predominantly nocturnal to more diurnal activity under the lower levels of light at night. Luarte et al. (2016) saw a similar switch in the sandhopper *Orchestoidea tuberculata* after seven days of observation, and shore crabs have been known to become less photonegative over time (Warman et al., 1993). Due to oxygen limitations, we could not extend the length of the assays to determine if this switch became more pronounced over time, but these results indicate that some crustaceans possess a certain level of resilience with regards to their relationship with light (Forward et al., 2007). As stated with *G. pulex*, whether this versatility is truly an adaptive advantage or not remains to be seen. There is evidence that amphipods can recover from short-term exposure to ALAN (Lynn et al., 2021), but long-term light pollution would be a more relevant condition to study due to permanent coastal light sources.

These results also do not incorporate the potential influence of tidal cycles on *E. marinus* behaviour. Future studies may address how ALAN impacts amphipod behaviour during and after tidal entrainment. It is likely that exposure to ALAN would be gated by tidal cycles due to the light scattering and absorbing effects of sea water. Moreover, as described below, behavioural control has been found to exhibit altered light sensitivity when it is dominated by circatidal versus circadian timekeeping. Thus, ALAN impacts in the intertidal zone will likely be modulated from those in the absence of tidal entrainment. Intertidal organisms demonstrate clear rhythmicity in terms of their movement (Chabot et al., 2016), feeding (Cheeseman et al., 2017), and reproduction (Barlow et al., 1986) related to tidal regimes and several studies have highlighted that this circatidal clock operates independently of the circadian clock (Satoh et al., 2008; Zhang et al., 2013). Following entrainment to tidal cycles, the talitrid amphipod *Parhyale hawaiensis* displayed peaks in activity correlated with high tide, regardless of daily light entrainment (Kwiatkowski et al., 2023). However, entrainment to daily light/dark cycles in the absence of tidal entrainment resulted in circadian rather than circatidal behaviour in this species. Under the latter circumstances. circadian rhythmicity was more pronounced in LL versus DD in *P. hawaiensis*, which matches our observations regarding *E. marinus*.

Other environmental stressors can influence gammarid behaviour as well. *E. marinus* and *G. pulex* both act as intermediate hosts to behavioural-altering parasites, the prevalence of which can reach up to 70%, depending on location and time of year (Bakker et al., 1997; Dudinák and Spakulová, 2003; Guler et al., 2015; Hohenadler et al., 2018). In addition, they inhabit areas which receive high levels of human pharmaceuticals via sewage effluent, with concentrations up to hundreds of micrograms per litre (Biel-Maeso et al., 2018; Daughton and Ternes, 1999; Kümmerer, 2010). While these two species are typically photophobic, both parasitism and exposure to anti-depressant medications can reverse their relationship with light by modifying serotonergic activity in the brain (Bakker et al., 1997; Guler et al., 2015; Helluy and Holmes, 1990). The compounding impacts of these behavioural-altering stressors under increasing levels of light pollution could have serious impacts on gammarid survival as predation risk increases, particularly if the hunting efficiency of predators is enhanced by ALAN.

While our results demonstrate significant behavioural changes due to light pollution, there is a high potential for ALAN to impact physiological processes as well. For example, photoperiod is a common factor in sex determination for invertebrates (Korpelainen, 1990), including *E. marinus*, with longer days yielding higher rates of male broods during the summer months (Guler et al., 2012). By obscuring the transition from day to night, anthropogenic light artificially extends the natural photoperiod and disrupts seasonal cycles in daylength. There is currently no information on how this could influence sex ratios at the population level over extended periods of time. Furthermore, disturbances to biological rhythms can lead to chronic stress (Ayalon et al., 2019; Levy et al., 2020), the physiological effects of which include altered metabolic function (Manríquez et al., 2019), tissue damage (Levy et al., 2020), and lowered immune function (Ouyang et al., 2017). Measuring concentrations of hemocyanin, an oxygen-transport protein in the hemolymph of invertebrates (Decleir and Richard, 1970), has successfully been used to monitor stress as a result of ALAN in talitrid amphipods (Lynn et al., 2022) and may prove likewise suitable for gammarids.

Given their significant ecological role, it is important to understand the long-term impacts of ALAN on invertebrate behaviour and how this will interact with other environmental stressors, such as those considered above as well as the warming or acidification of the oceans due to climate change. Along with light, temperature is another important zeitgeber for the circadian clock (Boothroyd et al., 2007; Currie et al., 2009; Glaser and Stanewsky, 2005) but, to our knowledge, there is no information available on the compounding impacts of climate change and light pollution on animal clocks, the disruption of which can carry significant consequences to an organism’s fitness. Considering the confluence of anthropogenic stressors affecting the environment, more comprehensive studies examining the compounding impacts of said stressors should be considered. Our findings, along with studies that used even lower lux levels (<1 lux), assert that any amount of light at night that sufficiently masks the natural light-dark cycle can dramatically alter animal behaviour and physiology, resulting in less activity (Latchem et al., 2021); altered foraging patterns (Gomes et al., 2021); reduced melatonin production (Franziska et al., 2020); and detrimental changes to reproductive success (Bird and Parker, 2014). While higher levels of light pollution have also been linked to avoidant behaviour (Abeel et al., 2016; Kobak and Nowacki, 2007); increased rates of aggression (Fonken et al., 2012); and significant shifts in predator-prey dynamics (Rydell, 1992; Yurk and Trites, 2000). The breadth of behavioural and physiological consequences as a result of light pollution at any scale encourages more extensive research that focuses on how individual changes in fitness may be carried across generations, leading to long-term repercussions for population and ecosystem health.

## Conclusions

Light pollution is an ever-expanding environmental hazard in an increasingly urbanised world. This study illustrates that artificial light at night can have strong but disparate effects on behavioural rhythms across closely related species. These effects may be further exacerbated by other anthropogenic stressors and carry significant physiological risks that have yet to be explored. Given the evolutionary significance of natural light cycles in guiding a variety of biological processes, it is essential to understand how light pollution may shape coastal and riparian ecosystems in the future.

## Supporting information

Raw data

## Acknowledgements

This work was supported by the INSPIRE Doctoral Training Programme and the Natural Environment Research Council (grant number NE/S007210/1). Thanks to the Invertebrate Facility of the School of Biological Sciences at the University of Southampton for providing space and equipment for the experiments that were conducted in this study. SCR was partially funded by Research England’s Expanding Excellence in England (E3) Fund.

