## Supplemental material for "Behavioural rhythms of two gammarid species *Echinogammarus marinus* and *Gammarus pulex* under increasing levels of light at night"

Table S1 Behavioural parameters as in Table 2 of main manuscript but separated for female (f) and male (m) *E. marinus* and *G. pulex.*

| **Species** | **Light regime** | **n_tot_** | | **n_al_** | | **n_act_** | | **%_al_** | | **%_act_** | | **Rhythmicity** | | | | **Total activity counts** | | | | **Nocturnality** | | | | **Period (hours)** | | | |
| --- | --- | --- | --- | --- | --- | --- | --- | --- | --- | --- | --- | --- | --- | --- | --- | --- | --- | --- | --- | --- | --- | --- | --- | --- | --- | --- | --- |
|  |  | **f** | **m** | **f** | **m** | **f** | **m** | **f** | **m** | **f** | **m** | **Avg_f_** | **± SE** | **Avg_m_** | **± SE** | **Avg_f_** | **± SE** | **Avg_m_** | **± SE** | **Avg_f_** | **± SE** | **Avg_m_** | **± SE** | **Avg_f_** | **± SE** | **Avg_m_** | **± SE** |
| *E. marinus* | LD | 15 | 17 | 15 | 17 | 15 | 17 | 100.0 | 100.0 | 100.0 | 100.0 | 2.19 | 0.15 | 2.54 | 0.23 | 2590 | 318 | 2640 | 370 | 0.73 | 0.03 | 0.75 | 0.03 | 24.0 | 0.05 | 23.7 | 0.32 |
|  | LA01 | 16 | 16 | 15 | 15 | 15 | 14 | 100.0 | 93.8 | 100.0 | 93.3 | 1.64 | 0.29 | 1.36 | 0.10 | 1964 | 424 | 1206 | 191 | 0.43 | 0.04 | 0.44 | 0.05 | 23.5 | 0.50 | 24.9 | 0.37 |
|  | LA05 | 13 | 19 | 10 | 11 | 10 | 10 | 76.9 | 57.9 | 100.0 | 90.1 | 1.31 | 0.19 | 1.39 | 0.14 | 1091 | 595 | 618 | 87 | 0.47 | 0.05 | 0.52 | 0.05 | 24.8 | 0.31 | 23.1 | 0.97 |
|  | LA30 | 16 | 16 | 16 | 14 | 15 | 14 | 100 | 87.5 | 93.8 | 100.0 | 1.36 | 0.20 | 1.36 | 0.12 | 2293 | 1244 | 730 | 115 | 0.43 | 0.04 | 0.45 | 0.03 | 23.9 | 0.33 | 23.9 | 0.07 |
|  | LA50 | 16 | 16 | 13 | 13 | 12 | 12 | 81.3 | 81.3 | 92.3 | 92.3 | 0.86 | 0.10 | 0.99 | 0.16 | 884 | 105 | 1195 | 587 | 0.43 | 0.03 | 0.46 | 0.05 | 26.4 | 2.06 | 24.3 | 0.14 |
|  | LA80 | 17 | 15 | 15* | 8* | 14 | 8 | 88.2 | 53.3 | 93.3 | 100.0 | 1.49 | 0.10 | 1.75 | 0.25 | 736 | 108 | 869 | 271 | 0.66 | 0.04 | 0.63 | 0.09 | 23..5 | 0.35 | 25.5 | 1.25 |
|  | LL | 16 | 16 | 16 | 15 | 8* | 14* | 100 | 93.8 | 50.0 | 93.3 | 0.99** | 0.08 | 1.72** | 0.23 | 424*** | 117 | 1548*** | 414 | 0.63 | 0.06 | 0.68 | 0.04 | 29.3 | 2.77 | 23.4 | 0.56 |
|  | DD | 32 | 32 | 16 | 18 | 14 | 14 | 50.0 | 56.3 | 87.5 | 77.8 | 0.90 | 0.07 | 0.96 | 0.04 | 1206 | 667 | 896 | 377 | 0.49 | 0.06 | 0.46 | 0.04 | 24.9 | 1.30 | 24.0 | 1.04 |
| *G. pulex* | LD | 16 | 16 | 15 | 15 | 8* | 14* | 93.8 | 93.8 | 53.3 | 93.3 | 1.16 | 0.13 | 1.32 | 0.18 | 945 | 314 | 1442 | 231 | 0.46 | 0.09 | 0.59 | 0.04 | 25.0 | 1.01 | 25.4 | 1.59 |
|  | LA01 | 16 | 16 | 15 | 15 | 12 | 15 | 93.8 | 93.8 | 80.0 | 100.0 | 0.79 | 0.08 | 0.95 | 0.06 | 1229** | 281 | 2567** | 487 | 0.45 | 0.04 | 0.53 | 0.03 | 24.5 | 4.77 | 24.5 | 0.55 |
|  | LA05 | 16 | 16 | 13 | 16 | 11 | 16 | 81.3 | 100.0 | 84.6 | 100.0 | 1.00 | 0.10 | 1.02 | 0.10 | 1712 | 499 | 2156 | 400 | 0.48 | 0.05 | 0.49 | 0.04 | 25.4 | 1.63 | 26.4 | 1.24 |
|  | LA30 | 17 | 15 | 17 | 15 | 9* | 13* | 100.0 | 100.0 | 52.9 | 86.7 | 0.94** | 0.16 | 1.46** | 0.14 | 673 | 157 | 1492 | 522 | 0.48 | 0.07 | 0.45 | 0.05 | 29.5 | 5.50 | 24.8 | 1.43 |
|  | LA50 | 16 | 16 | 13 | 13 | 8 | 13 | 81.3 | 81.3 | 61.5 | 100.0 | 0.83 | 0.13 | 0.97 | 0.06 | 907 | 263 | 846 | 151 | 0.52 | 0.06 | 0.47 | 0.04 | 23.5 | 2.18 | 26.4 | 1.32 |
|  | LA80 | 16 | 16 | 12 | 15 | 10 | 13 | 75.0 | 93.8 | 83.3 | 86.7 | 0.96 | 0.14 | 1.24 | 0.14 | 746 | 116 | 1501 | 326 | 0.43 | 0.05 | 0.36 | 0.04 | 24.2 | 0.44 | 24.6 | 0.39 |
|  | LL | 16 | 15 | 15 | 13 | 12 | 10 | 93.8 | 86.7 | 80.0 | 76.9 | 0.89 | 0.06 | 0.86 | 0.12 | 1010 | 157 | 1468 | 431 | 0.56 | 0.05 | 0.53 | 0.04 | 23.3 | 2.68 | 27.0 | 2.33 |
|  | DD | 16 | 16 | 16 | 15 | 15 | 15 | 100 | 93.8 | 93.8 | 100.0 | 0.79 | 0.06 | 0.79 | 0.05 | 1640* | 420 | 2892* | 1894 | 0.46 | 0.02 | 0.48 | 0.01 | 25.0 | 1.00 | 24.8 | 3.25 |
| ntot = total number of individuals at the beginning of the assays; nal = # of individuals alive at the end of the eight day assays; %al = (nal /ntot)·100; nact = # of active individuals with total counts ≥ 200; %act = (nact/nal)·100; Avg = average; SE = standard error; %SR = % active individuals showing strongly rhythmic behaviour (RRP ≥ 1.5); %WR = % active individuals showing weakly rhythmic behaviour (RRP = 1–1.49); %AR = % active individuals showing arhythmic behaviour (RRP < 1). Different superscript letters denote significant differences (p < 0.05) between treatments from the Fisher’s exact tests and the Mann-Whitney U-tests. *E. marinus* and *G. pulex* were analysed separately. | | | | | | | | | | | | | | | | | | | | | | | | | | | |

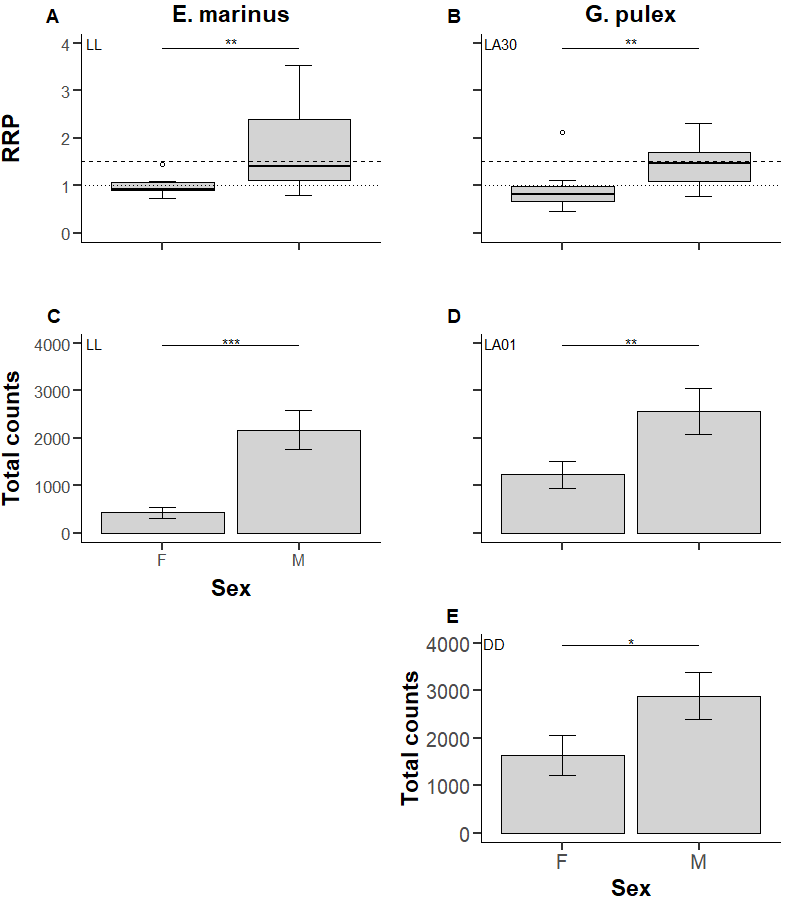

Figure S1 Activity of female and male *E. marinus* (left column) and *G. pulex* (right column). Boxplots of relative rhythmic power (RRP; A-B) - how strongly the activity repeated over a set period (e.g. 24 hours). Dotted line = 1 (rhythmic); Dashed line = 1.5 (strongly rhythmic). Barplots of total counts (C-E) - total amount of activity counts logged during the seven-day assay period. Assays occurred over seven days with 3-hour ramping between light transitions. Asterisks denote significance levels from the Mann-Whitney U-tests (0.5*0.1**0.001***0.0001). Light conditions outlined in Table 1 of main manuscript.

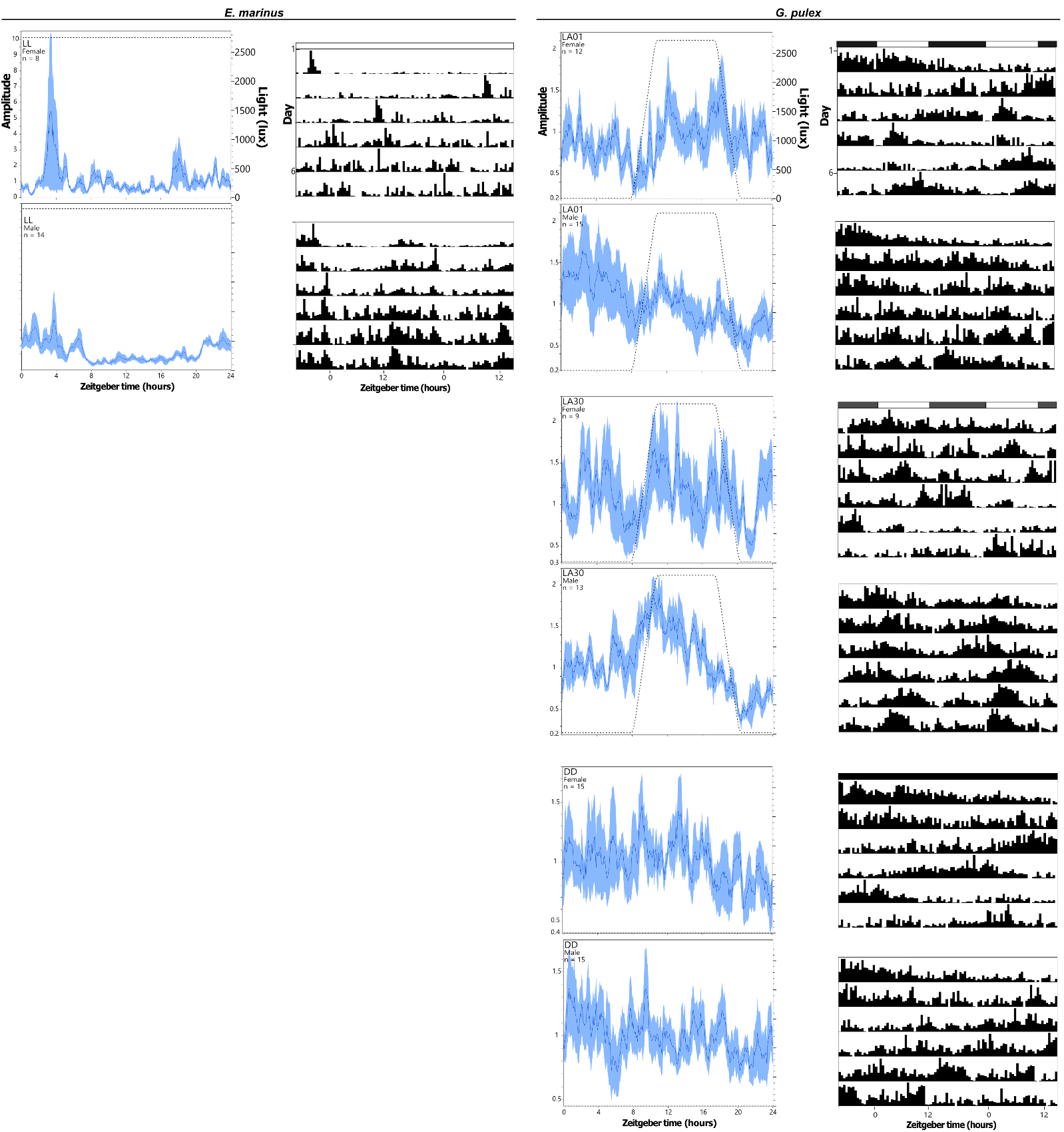

Figure S2 Average activity profiles (columns 1 & 3) and double-plotted actograms (col. 2 & 4) of female and male *E. marinus* (col. 1 & 2) and *G. pulex* (col. 3 & 4). Activity profiles display average activity levels (black line) with standard deviation (grey areas), along with the light levels (dotted lines) over 24 hours. Actograms display the average, normalised behaviour across the 7-day assays, with each row showing two 24-hour cycles. Bars above actograms denote light levels; on/off transitions are not shown. Y axes are the same within columns. Light conditions outlined in Table 1 of main manuscript.
